# Towards improved resistance of *Corynebacterium glutamicum* against nisin

**DOI:** 10.1101/2021.08.09.454123

**Authors:** Dominik Weixler, Oliver Goldbeck, Gerd. M. Seibold, Bernhard J. Eikmanns, Christian U. Riedel

**Author notes:** Corresponding author. Institute of Microbiology and Biotechnology, University of Ulm, Albert-Einstein-Allee 11, 89081 Ulm, Germany.

## Abstract

The bacteriocin nisin is one of the best studied antimicrobial peptides. It is widely used as a food preservative due to its antimicrobial activity against various Gram-positive bacteria including human pathogens such as *Listeri*a *monocytogenes* and others. The receptor of nisin is the universal cell wall precursor lipid II, which is present in all bacteria. Thus, nisin has a broad spectrum of target organisms. Consequently, heterologous production of nisin with biotechnological relevant organisms including *Corynebacterium glutamicum* is difficult. Nevertheless, bacteria have evolved several mechanisms of resistance against nisin and other cationic antimicrobial peptides (CAMPs). Here, we transferred resistance mechanisms described in other organisms to *C. glutamicum* with the aim to improve nisin resistance. The presented approaches included: expression of (i) nisin immunity genes *nisI* and/or *nisFEG* or (ii) nisin ABC-transporter genes of *Staphylococcus aureus* and its homologues of *C. glutamicum*, (iii) genes coding for enzymes for alanylation or lysinylation of the cell envelope to introduce positive charges, and/or (iv) deletion of genes for porins of the outer membrane. None of the attempts alone increased resistance of *C. glutamicum* more than two-fold. To increase resistance of *C. glutamicum* to levels that will allow heterologous production of active nisin at relevant titers, further studies are needed.

## 1 Introduction

Bacteriocins are ribosomally synthesized peptides naturally produced by various bacteria with antimicrobial activity against a broad range of bacteria (Cotter et al., 2013; Meade et al., 2020). One of the best characterized bacteriocin is nisin produced by *Lactococcus lactis* species (Lubelski et al., 2008). It consists of 34 amino acids and belongs to the group of class Ia bacteriocins also termed lantibiotics based on their (methyl-)lanthionine rings, which are introduced during posttranslational modification (Arnison et al., 2013). Nisin shows high antimicrobial activity against several Gram-positive bacteria including the human pathogens *Staphylococcus aureus* (Brumfitt et al., 2002; Jensen et al., 2020) and *Listeria monocytogenes* (Benkerroum and Sandine, 1988; Harris et al., 1991).

Nisin is synthesized as inactive pre-peptide (prenisin) in the cytoplasm, exported to the extracellular space and activated by the nisin protease NisP that cleaves off the leader peptide (Lubelski et al., 2008; Xu et al., 2014). Like other antimicrobial peptides, nisin has a positively charged N-terminus that facilitates interaction with the negatively charged envelope of target organisms. The specific receptor of nisin is the universal cell wall precursor molecule lipid II (Breukink et al., 1999; Brötz et al., 1998), which is located in the outer leaflet of the cell membrane.

The antimicrobial activity of nisin is based on a dual mode of action. On the one hand, by binding to lipid II (Hsu et al., 2004), nisin prevents incorporation of this cell wall precursor into the nascent peptidoglycan chain and thus inhibits cell wall biosynthesis and growth (Kuipers et al., 2006). Additionally, nisin and lipid II are able to assemble to pore-forming complexes consisting of 8 molecules of nisin and 4 molecules lipid II (Hasper et al., 2004) that mediate killing of target bacteria (Breukink et al., 1999).

Nisin was approved by the FDA and received “generally regarded as safe” status (Delves-Broughton, 1996). Currently, nisin is produced with optimized LAB strains on complex milk- or whey-based substrates resulting in low product purity and/or elaborate and expensive downstream processing (de Arauz et al., 2009; Juturu and Wu, 2018; Li et al., 2002).

Recombinant nisin production using biotechnological workhorse organisms on an industrial scale has not been demonstrated so far. A potential candidate for recombinant nisin production is *Corynebacterium glutamicum*. This organism is well established in biotechnological processes and was shown to be a suitable production host for a variety of different compounds including bulk chemicals, L-amino acids or therapeutic proteins such as single chain antibody fragments (Becker et al., 2018; Wolf et al., 2021; Yim et al., 2016, 2014). Recently, recombinant synthesis of the class IIa bacteriocin pediocin PA1 using *C. glutamicum* was successfully demonstrated (Goldbeck et al., 2021, submitted). However, while *C. glutamicum* is resistant to pediocin PA1, nisin targets the peptidoglycan precursor lipid II found in all bacteria. Consequently, nisin has a wide spectrum of target organisms including *C. glutamicum* (Goldbeck et al., 2021, submitted). Thus, production of active nisin using normal production strains of *C. glutamicum* will be difficult.

In addition to the natural producers, which require efficient protection against the bactericidal activity of nisin, several other organisms have evolved mechanisms of resistance against nisin. This includes general mechanisms of resistance to cell envelope stress including cell wall thickening or cell envelope alterations like incorporation of positively charged substituents to alter the overall negative charge of the surface (Draper et al., 2015). A well characterized example is D-alanylation of teichoic acids catalyzed by the *dlt* operon (Neuhaus and Baddiley, 2003) that was discovered in different Firmicutes and demonstrated to confer resistance against a broad range of CAMPs (Abi Khattar et al., 2009; Kovács et al., 2006; McBride and Sonenshein, 2011a; Peschel et al., 1999). Another modification altering surface charge is lysinylation of the membrane phospholipid phophatidylglycerol (PG) by e.g. the multi-resistance-factor F (MprF) found in different Firmicutes like *S. aureus* (Ernst and Peschel, 2011) or *L. monocytogenes* (Thedieck et al., 2006). Loss of MprF function increases sensitivity of *S. aureus* against various AMPs (Ernst et al., 2009; Nishi et al., 2004). A similar effect is described for the bifunctional lysine-tRNA ligase/phosphatidylglycerol lysyltransferase (LysX) of *Mycobacterium tuberculosis* that contributes to an increased resistance against CAMPs (Maloney et al., 2009).

More specific resistance mechanisms are often based on ABC-transporter systems and their regulatory two or multiple-component systems (Draper et al., 2015). In general, ABC-transporters involved in CAMP resistance can be grouped in two classes (Clemens et al., 2018). One group is the bacitracin efflux ABC-transporter (BceAB-type), first characterized in *B. subtilis* (Ohki et al., 2003). Further prominent members are the VraDE transporter from *S. aureus* (Hiron et al., 2011) or NsrFP of *S. agalactiae* (Reiners et al., 2017). Both were shown to confer resistance to nisin, bacitracin or other CAMPs (Arii et al., 2019; Hiron et al., 2011; Zaschke-Kriesche et al., 2020). The other group consists of cationic peptide resistance ABC-transporters (CrpABC-type) including the well characterized nisin immunity system NisFEG from *L. lactis* (Alkhatib et al., 2014) or CprABC of *Clostridioides difficile* strains (McBride and Sonenshein, 2011b). These transporters are mostly found in natural AMP producers, are highly specific, and confer immunity against their own product. Moreover, CrpABC-type transporters work cooperatively with additional immunity systems. For example, NisFEG of *L. lactis* nisin producers acts together with the lipoprotein NisI to achieve full immunity against nisin (Stein et al., 2003).

In the present study, we tried to increase resistance of *C. glutamicum* to nisin by introducing genes for several of the described resistance or immunity mechanisms or deletion of porins in the mycolic acid. The aim was to render this biotechnological platform organism resistant to a degree that would allow recombinant production of active nisin.

## 2 Materials and Methods

### 2.1 Bacterial strains and growth conditions

All strains and plasmids used in this study are listed in **Supplementary Table S1**. *C. glutamicum* and *E. coli* were cultivated in 2xTY complex medium and incubated with agitation at 30 °C and 37 °C, respectively. *B. subtilis* and *S. aureus* were cultivated on solid Brain Heart Infusion (BHI) medium containing agar (15 g/l). *L. lactis* subsp. *lactis* B1629 was cultivated on solid GM17 medium. For induction of gene expression, isopropyl-β-D-thiogalactoside (IPTG) and/or anhydrotetracycline (aTc) was used in the indicated concentrations. For selection, kanamycin (25 µg/ml) was used if appropriate.

### 2.2 Cloning procedures

For molecular cloning procedures standard reagents were used according to the manufacturer’s instructions. PCR reactions were performed in a C100 thermocycler (Bio-Rad Laboratories, Munich, Germany), nucleotides were purchased from Bio-Budget (Krefeld, Germany). Genes for nisin immunity proteins *nisF nis E nisG* and *nisI* were amplified using genomic DNA of the natural nisin producer strain *L. lactis* subsp. *lactis* B1629. In addition, a *nisI* gene with codon-usage optimized for expression in *C. glutamicum* (*nisI*^*CO*^) was obtained as a synthesized DNA fragment from a commercial service provider (Eurofins genomics; Ebersberg, Germany). The ABC-transporter genes *vraD* and *vraE* were amplified from *S. aureus* MRSA ATCC 43300. Genes *cg2812-11, cg3322-20* and *cg1103* were amplified from *C. glutamicum* CR099. All primers and the codon-optimized gene sequence of *nisI*^*CO*^ are listed in **Supplementary Table S2**. All genes were amplified using primers containing ribosomal binding sites for *C. glutamicum* and overlapping regions for fusion in the correct order into the indicated expression vectors by Gibson assembly (Gibson et al., 2009). Generally, competent *E. coli* DH5α cells as the cloning host. Prior to transformation into the respective target organisms, all plasmids were isolated from *E. coli* and correct cloning was verified by restriction analysis and Sanger sequencing (Eurofins genomics). *C. glutamicum* CR099 was transformed by electroporation as described previously (Tauch et al., 2002).

### 2.3 Construction of deletion mutants

Deletion mutants were constructed based on a previously described CRISPR-Cpf1 method for genome editing in *C. glutamicum* (Jiang et al., 2017). The vector backbone of pJYS3_Δ*crtYf* (Jiang et al., 2017) was amplified by inverse PCR to remove *crtY*-specific sequences and subsequently ligated with a non-coding 500 bp DNA fragment introducing unique *Sma*I, *Swa*I, and *BamH*I restriction sites and yielding plasmid pJYS3-KH. Appropriate protospacer sequences including TTTN protospacer-adjacent motifs (PAM) for guidance of the Cpf1 enzyme were defined in the genes to be deleted (**Supplementary Table S3**). Potential off-targeting was checked via BLAST analyses of the selected adjacent protospacer regions against the genome of *C. glutamicum* ATCC13032. Possible secondary structures or hairpin formation was evaluated by the online software oligo calc (Kibbe, 2007). Corresponding sgRNA genes of 108 bp including the promotor sequence, 20 bp overlap handle and *BamH*I/*Swa*I restriction sites were obtained by overlap extension PCR using two oligonucleotides as template and sub-cloned into pJet1.2 blunt vector (Thermo Scientific) according to the manufacturer’s protocol. These sgRNA fragments and pJYS3-KH were cut using restriction enzymes *BamH*I and *Swa*I and ligated by T4-DNA ligase (Thermo Scientific) resulting in plasmids pJYS3_*sgRNAx*. For homologous recombination, each 500 bp up- and downstream of the desired gene were designed and cloned into a *Sma*I linearized pJYS3_*sgRNAx* vector via Gibson assembly. *C. glutamicum* CR099 was transformed with the assembled deletion plasmids and plated on 2x TY agar plates containing kanamycin for selection. After 3 days of incubation at 30 °C positive clones were purified by re-streaking on selective agar for three passages and obtained deletion mutants were then cultivated overnight at 34 °C in 2x TY medium w/o kanamycin for plasmid curing. Afterwards strains were checked by screening for absence of growth on agar plates containing kanamycin and absence of PCR product specific for the plasmid. Mutants were finally checked by sequencing of the PCR product obtained from amplification of the distinct genome region.

### 2.4 Growth inhibition assay

To assess resistance to AMPs, a growth inhibition assays were performed in micro titer plate format as described previously (Wiegand et al., 2008) with minor modifications. Serial 1:2 dilutions of a standard solution of nisin Z (Sigma-Aldrich) were prepared in 2xTY and 100 µl were distributed in 96-well plates (Sarstedt, Nümbrecht, Germany). Bacteria of the test strains were inoculated from agar plates and cultivated overnight in 2xTY medium containing kanamycin. IPTG (0.1 mM) and/or ATC (0.25 µg/ml) were added to induce gene expression where appropriate. o/N cultures were diluted in fresh medium containing the inducers and 100 µl were added to each well of the assay plate to a final optical density at 600 nm (OD_600_) of 0.05 in the assay. Assay plates were incubated with agitation at 30 °C. Growth was determined after 24 h by measuring OD_600_ in an Infinite M100 plate reader (Tecan, Männedorf, Suisse). The minimal inhibitory concentration (MIC) was calculated based on the lowest dilution of the nisin standard.

## 3 Results and Discussion

Currently, biotechnological nisin production is exclusively performed with natural producer strains in batch fermentation processes on complex milk- or whey-based substrates (de Arauz et al., 2009; Juturu and Wu, 2018; Li et al., 2002). Complex media result in difficult and expensive downstream processing and purification (Abbasiliasi et al., 2017) and consequently, nisin is marketed only as partially purified preparation containing max. 2.5% of active product (de Arauz et al., 2009; Taylor et al., 2007). Production with industrial workhorse organisms such as *C. glutamicum* on defined minimal media containing sustainable substrates may result in more efficient and economic process and higher product purities.

A major challenge towards heterologous nisin production using *C. glutamicum* is low resistance of the production host against the product. In a previous study, *C. glutamicum* was described to grow in the presence of nisin at concentrations of 40 µg/ml of nisin albeit at reduced growth rates (Sieger et al., 2013). In a recent study, we observed no growth after 24h at 0.5-1 µg/ml of nisin and slow but measurable growth at concentrations below 0.5 µg/ml (Goldbeck et al., 2021, submitted). These differences are related to different methods to calculate nisin concentrations. Our own calculations are based on the mass of active compound in the commercial product (2.5%). When calculating concentrations using the total mass of the commercial nisin preparation this would result in a concentration of approx. 20 µg/ml at which growth is still observed, which is close to the value published by others (Sieger et al., 2013). In any case, inhibition of growth of the producing organism by product concentrations of 0.5 µg/ml is too low for an efficient process. In the present study, we thus tried to improve nisin resistance of *C. glutamicum* by generation of recombinant strains expressing genes known to confer nisin/CAMP resistance in other Gram-positive bacteria. In order to avoid degradation of the product, we excluded resistance mechanisms that are based on specific proteolytic degradation of nisin described e.g. in *Streptococcus agalactiae* (Khosa et al., 2013).

### 3.1 Immunity protein and ABC transporter of natural producer L. lactis

An obvious approach to increase resistance against AMPs is to introduce the corresponding immunity systems from in natural producers. For nisin, two immunity systems are encoded in the nisin operon of *L. lactis* producer strains. The first one is the lipoprotein NisI, which specifically binds nisin thereby preventing or reducing binding to its target lipid II (AlKhatib et al., 2014; Koponen et al., 2004; Stein et al., 2003; Takala and Saris, 2006). The second immunity system is the ABC-transporter NisFEG that expels nisin from the cytoplasmic membrane (Alkhatib et al., 2014). The genes (*nisI* and *nisFEG*) were amplified from genomic DNA of the natural nisin Z producer *L. lactis* subsp. *lactis* B1629 and cloned into the *C. glutamicum* expression vector pEKEx2 under the control of the P_*tac*_ promotor.

Recombinant *C. glutamicum* strains in the CR099 background (Baumgart et al., 2013) harboring these plasmids were analyzed for nisin resistance with expression induced by IPTG (**Figure 1**). A marginal improvement in resistance could be observed with expression of *nisFEG* compared to the empty vector control strain could be observed at a nisin concentration of 1562 ng/ml as indicated by an OD_600_ slightly above completely inhibited cultures. By contrast, expression of *nisI* had an adverse effect on resistance with OD_600_ markedly lower at 781 ng/ml nisin and below. Further experiments with *C. glutamicum* CR099/pEKEx-*nisI*^CO^, strain expressing a synthetic *nisI* gene optimized for codon-usage of *C. glutamicum* confirmed, that expression of *nisI* does not increases resistance to nisin (data not shown). Here, no difference in final OD_600_ between *C. glutamicum* CR099/pEKEx-*nisI*^CO^ and the empty vector control strain were observed after 24 h at any of the concentrations tested. This suggests that the slightly reduced OD600 of *C. glutamicum* CR099/pEKEx-*nisI* may be caused by expression of a *L. lactis* gene with low G+C content of 34 %. For *L. lactis*, it was shown that full immunity against nisin requires of both NisI and NisFEG in combination (Ra et al., 1999; Stein et al., 2003). Thus, we constructed the strain *C. glutamicum* pOGOduet-*nisI*-*nisFEG* for dual expression of both systems in combination. However, combined expression of *nisI* and *nisFEG* also did not lead increased resistance of *C. glutamicum* CR099 to nisin (**Supplementary Figure S1**). In summary. expression of the resistance mechanisms of the native nisin Z producer *L. lactis* in *C. glutamicum* did not result in a considerable improvement of nisin resistance compared to the empty vector control. By contrast, heterologous expression of *nisI* and/or *nisFEG* in *B. subtilis* yielded strains with improved nisin resistance (Hansen et al., 2009). Both *B. subtilis* and *L. lactis* belong to the phylum Firmicutes and thus share similarities regarding the composition of membrane and cell wall. Furthermore, *B. subtilis* isolates are able to synthesize subtilin, a lantibiotic related to nisin (Paik et al., 1998; Rintala et al., 1993). *C. glutamicum*, on the other hand, belongs to the actinobacteria that are characterized by an unique cell wall composition including the additional (outer) mycolic acid membrane (Burkovski, 2013; Marchand et al., 2012). Thus, an appropriate cell membrane surrounding could be a crucial requirement for a full functionality of transmembrane and lipoproteins such as NisI and NisFEG.

**Figure 1:**
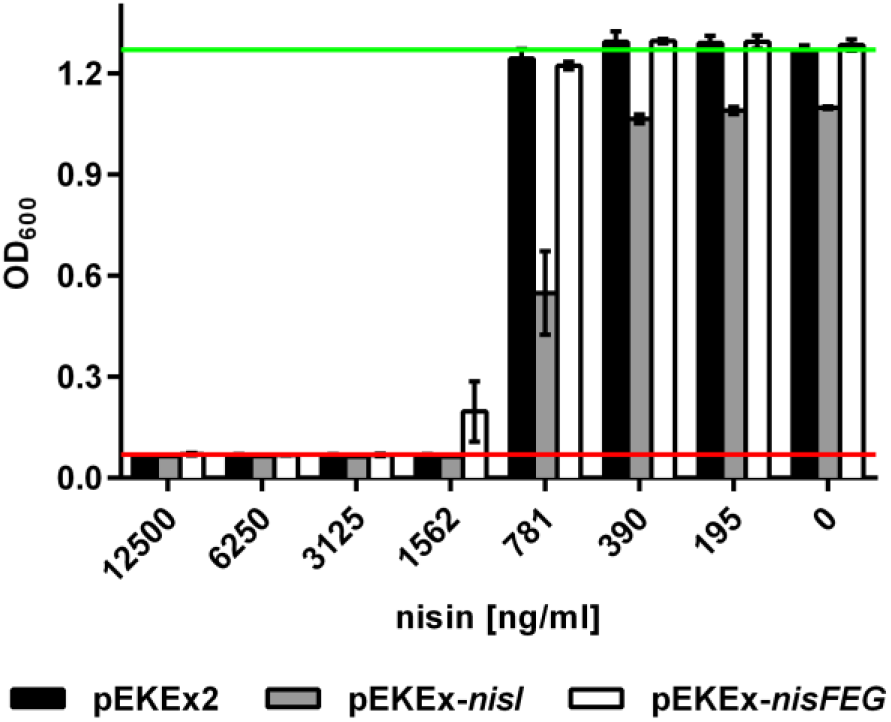
Resistance of *C. glutamicum* strains expressing nisin immunity genes of the native producer *L. lactis* B1629. Strains CR099/pEKEx-*nisI* (grey bars), CR099/pEKEx-*nisFEG* (white bars) or the empty vector control strain CR099/pEKEx2 (black bars) were cultivated in 2xTY medium in 96-well microtiter plates in the presence of nisin at the indicated concentrations. For induction of gene expression 0.1 mM IPTG was added. Optical density at 600 nm (OD_600_) was determined after 24 h of incubation. The red and green lines indicate OD_600_ of the positive (i.e. complete inhibition of growth) or negative (i.e. in the absence of nisin) control, respectively. All values are mean ± standard deviation of n = 3 cultivations of the test strain.

### 3.2 ABC transporters involved in CAMP resistance

In addition to the highly specific immunity systems of natural nisin producers, several mechanisms found in other organisms that increased resistance against CAMPs in general (Draper et al., 2015). A quite well characterized system is BceAB-type ABC-transporter VraDE involved in bacitracin and nisin detoxification (Arii et al., 2019; Hiron et al., 2011). A BLAST search of the *C. glutamicum* genome identified an ABC-transporter encoded by the genes *cg2812-11* as a potential VraDE homologues. In a previous study, *cg2812-11* were shown to be regulated by the three component system EsrISR (Kleine et al., 2017). This three component system is involved in the cell envelope stress response and its expression is increased in the presence of bacitracin (Kleine et al., 2017). Another operon showing increased expression upon bacitracin exposure comprises the genes *cg3322-20*, also encoding an ABC transporter. Deletion of either *esrISR* or *cg3322-20* resulted in increased sensitivity to bacitracin (Kleine et al., 2017). Thus, we generated *C. glutamicum* CR099 strains (over-)expressing genes *cg3322-20, cg2812-11* or *vraDE* individually, which were cloned into pEKEx2 under the control of the P_*tac*_ promotor. Analysis of the strains of these recombinant strains revealed that growth of most strains was completely abolished by nisin concentrations of 2.5 µg/ml and above (**Figure 2A**). For *C. glutamicum* CR099/pEKEx-*vraDE*, a slightly increased nisin resistance was observed.

**Figure 2:**
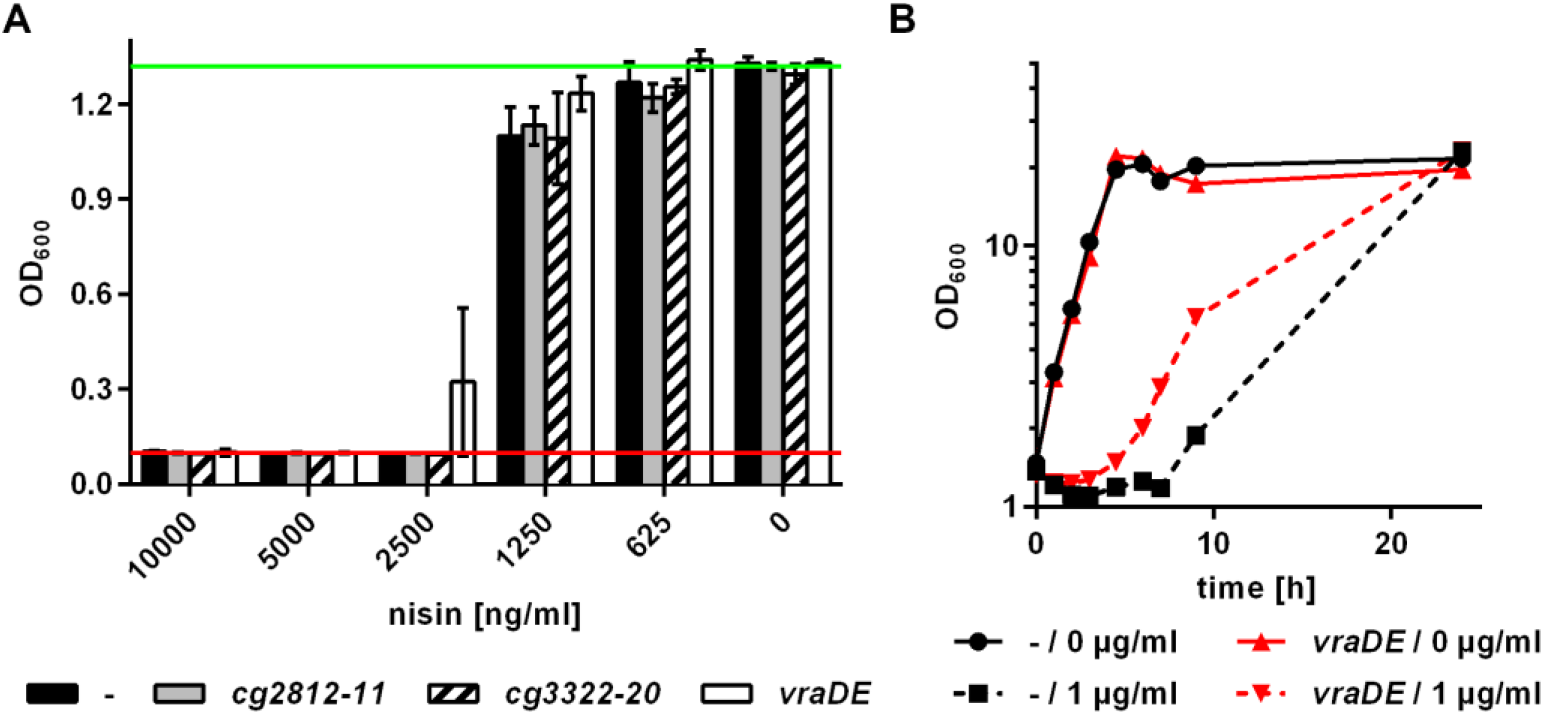
Resistance of *C. glutamicum* strains expressing genes for ABC transporters. (A) Strains CR099/pEKEx-*cg2812-11* (grey bars), CR099/pEKEx-*cg3322-20* (hatched bars), CR099/pEKEx-*vraDE* (white bars) or the empty vector control strain CR099/pEKEx2 (black bars) were cultivated in 96-well microtiter plates. OD_600_ was determined after 24 h of incubation. The red and green lines indicate OD_600_ of the positive (i.e. complete inhibition of growth) or negative (i.e. in the absence of nisin) control, respectively. Values are mean ± standard deviation of n = 3 cultivations of the test strain. (B) CR099/pEKEx-*vraDE* (red) or the empty vector control strain CR099/pEKEx2 (black) was grown in shake flasks in the presence (broken lines) or absence (solid lines) of nisin (1 µg/ml) and OD_600_ was determined at the indicated timepoints. In all experiments, 2xTY medium supplemented with 0.1 mM IPTG for induction of gene expression and nisin at the indicated concentrations was used.

To confirm this slight increase in resistance, we performed further experiments measuring growth in shake flasks in presence of nisin (1 µg/ml), i.e. below the MIC. This revealed that the extended lag phase caused by nisin was improved by expression of *vraDE*. Effects during lag phase are not captured by the microtiter plate assay. Further experiments need to be performed to clarify if similar effects may be observed for expression of *cg2812-11* or *cg3322-20* may have similar effects at lower nisin concentrations. (**Figure 2B**).

Recently, we performed an analysis of bacteriocin gene clusters and associated resistance mechanisms in the genus *Corynebacterium* (Goldbeck et al., 2021). Interestingly, several genomes of the genus harbored lantibiotic gene clusters and associated ABC-transporters and some strains were able to tolerate nisin concentrations of up to 25 µg/ml. However, expression of genes for these ABC transporters in *C. glutamicum* ATCC13032 also improved resistance to a maximum of 5 µg/ml of nisin. Overall, our results suggest that expression of genes for ABC-transporters with a reported or predicted role in CAMP resistance only has little effects on nisin resistance of *C. glutamicum* and best results are achieved with ABC transporters of other corynebacteria.

### 3.3 Cell membrane modification

A central mechanism of CAMP resistance found in several organisms is based on alteration of the overall cell membrane charge (Draper et al., 2015; Ernst and Peschel, 2011). Bacterial cell membranes are mainly composed of negatively charged phospholipids phosphatidyl glycerol, diphosphatidylglycerol or phosphatidylinositol (Sohlenkamp and Geiger, 2015). Similarly, the glycerol- and sugar-phosphate backbones of teichoic acids in the cell wall of Gram-positive bacteria has a negative net charge (Brown et al., 2013; Percy and Gründling, 2014). Consequently, CAMPs are attracted to cell wall and membrane where they are then able to bind to their receptor lipid II. To reduce electrostatic attraction of CAMPs to the cell surface, bacteria have evolved mechanisms for incorporation of positively charged residues into the (lipo)teichoic acids and phospholipids. One example is alanylation of teichoic acids by enzymes encoded the *dlt* operons found in e.g. *S. aureus, C. difficile* or *L. monocytogenes* (Abachin et al., 2002; McBride and Sonenshein, 2011a; Neuhaus and Baddiley, 2003; Peschel et al., 1999). The aminogroup of alanine residues carries a positive charge that is reported to confer a shielding effect by masking the negatively charged teichoic acids and repulsion of CAMPs (Neuhaus and Baddiley, 2003). *C. glutamicum* does not possess teichoic acids and thus the substrates for alanylation by the Dlt enzymes are missing.

Another mechanism that alters the overall negative charge of the cell membrane is the substitution of phosphatidylglycerol (PG) with positively amino acids like lysine (Draper et al., 2015; Ernst and Peschel, 2011). In *S. aureus*, lysinylation is carried out by the MprF protein (Ernst et al., 2009; Peschel et al., 2001). A similar reaction is carried out by the bifunctional lysine-tRNA ligase/phosphatidylglycerol lysyltransferase LysX in *M. tuberculosis* (Maloney et al., 2009). Thus we tried to express the *lysX* homologue *cg1103* (Smith et al., 2015) coding for an aminoacyl-phosphatidylglycerol synthase in *C. glutamicum*. However, no increase in nisin resistance was observed for the recombinant strain *C. glutamicum* CR099/pEKEx-*cg1103* (**Figure 3**). Similarly, we created the strain *C. glutamicum* CR099/pEKEx-*mrpF* that expresses of *mprF* from *B. subtilis*. However, this strain was characterized by a very long lag-phase, inconsistent growth behavior and genetic instability with plasmid loss (data not shown), and hence was not further analyzed for nisin resistance.

**Figure 3:**
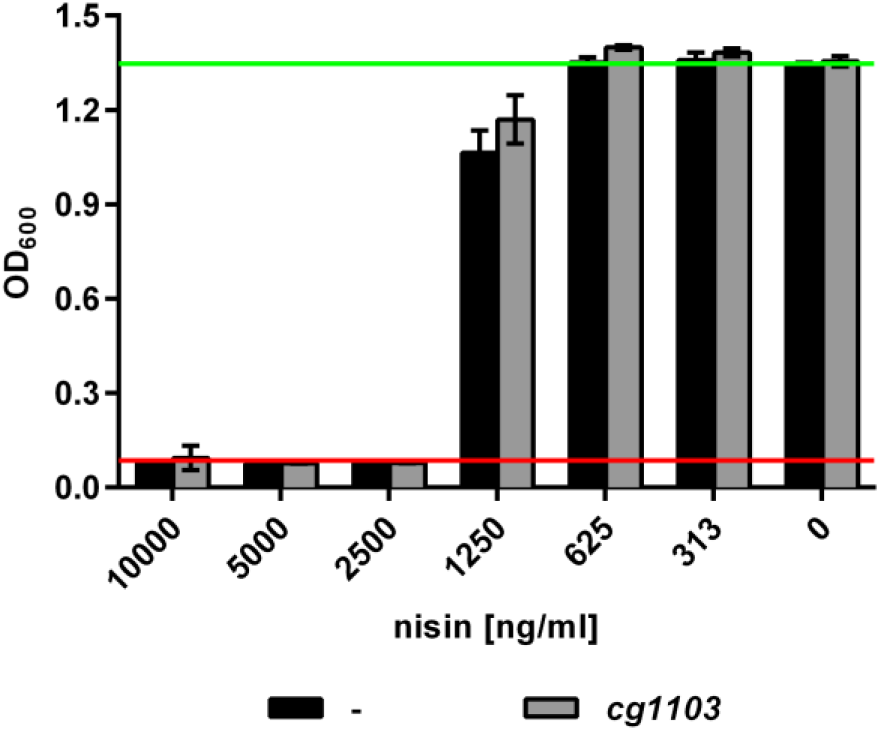
Nisin resistance of *C. glutamicum* strains expressing the *lysX* homologue *cg1103*. *C. glutamicum* CR099/pEKEx-*cg1103* (grey bars) or the empty vector control strain CR099/pEKEx2 (black bars) were cultivated in 96-well microtiter plates in 2xTY medium supplemented with 0.1 mM IPTG for induction of gene expression and nisin at the indicated concentrations. OD_600_ was determined after 24 h of incubation. The red and green lines indicate OD_600_ of the positive (i.e. complete inhibition of growth) or negative (i.e. in the absence of nisin) control, respectively. Values are mean ± standard deviation of n = 6 cultivations of the test strain.

Alternatively, we tried to achieve a shielding effect in *C. glutamicum* CR099 by adding CaCl_2_ to the cultivation medium. Following dissociation in solution, Ca^2+^ ions are assumed to bind to negatively charged components of the cell envelope (Thomas III and Rice, 2014). Addition of CaCl_2_ (2 g/l) to the medium improved nisin resistance approx. 2x fold and 5 µg/ml were required to completely inhibit growth of *C. glutamicum* CR099/pEKEx2 (**Figure 4**). As corynebacteria are lacking teichoic acids, this effect is most likely based on interactions of Ca^2+^ ions to negatively charged moieties of membrane lipids and cell wall glycopolymers containing phosphate groups such as arabinogalactans and lipoarabinomannans (Burkovski, 2013; Weidenmaier and Peschel, 2008).

**Figure 4:**
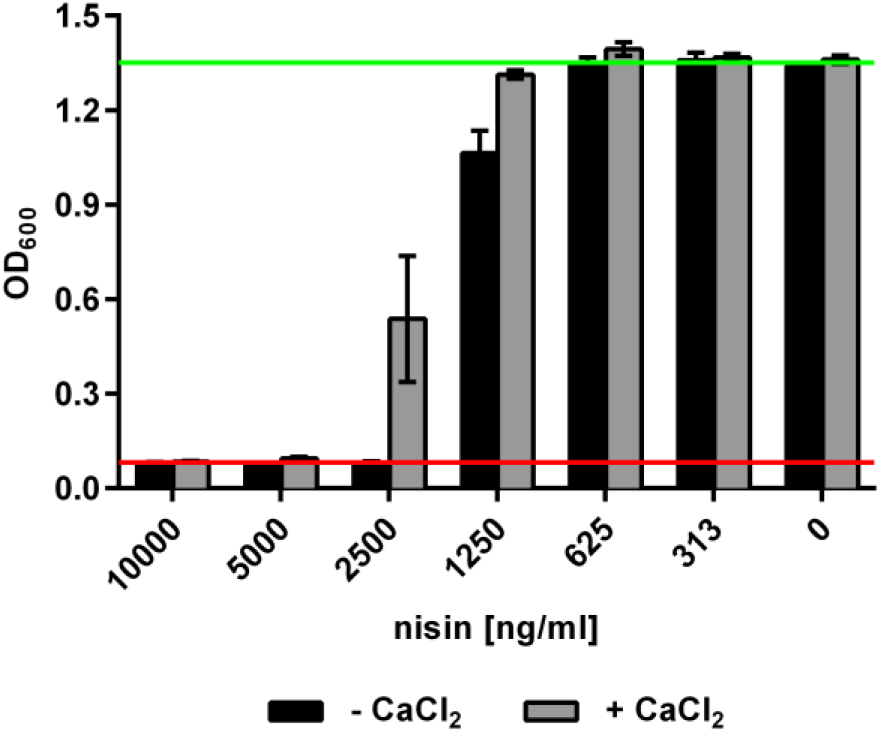
Effect of CaCl_2_ on resistance of *C. glutamicum* CR099 to nisin. Bacteria were grown in 96-well microtiter plates in 2xTY medium supplemented with 2 g/l CaCl_2_ and nisin at the indicated concentrations. OD_600_ was determined after 24 h of incubation. The red and green lines indicate OD_600_ of the positive (i.e. complete inhibition of growth) or negative (i.e. in the absence of nisin) control, respectively. Values are mean ± standard deviation of n = 6 cultivations of the test strain.

Overall, approaches that introduces positive charges or shields negative charges in the cell envelope appear to be able to increase resistance of *C. glutamicum* to nisin and other CAMPs to some extent.

### 3.4 Potential CAMP uptake systems

The cell envelope of actinobacteria including *C. glutamicum* is characterized by an additional layer covering the peptidoglycan cell wall that contains a mycolic acid membrane and resembles to some extent the outer membrane of Gram-negative bacteria (Burkovski, 2013; Puech et al., 2001). Similar to the outer membrane of Gram-negatives, the mycolic acid membranes of *C. glutamicum* and other actinobacteria harbor porins that facilitate passage of charged molecules (Costa-Riu et al., 2003a, 2003b; Hünten et al., 2005; Lichtinger et al., 1999, 1998; Ziegler et al., 2008). For *C. glutamicum*, four potential porins (PorA, PorH, PorB and PorC) are described that share structural similarities and are all substrates for posttranslational modification by mycoloylation (Issa et al., 2017). PorA and PorH were shown to form heteromeric channel complexes that allow translocation of positively charged macromolecules (Barth et al., 2010; Rath et al., 2011). Moreover, deletion of *porA* led to decreased sensitivity to positive charged antibiotics (Costa-Riu et al., 2003a). On the other hand, channels formed by PorB and PorC were shown to allow passage of negatively charged macromolecules (Costa-Riu et al., 2003b). To test if deletion of any of the porins may increase resistance to nisin by restricting access of this positively charged peptide to the membrane, we constructed a series of deletion mutants. Single deletion mutants lacking *porA* or *porH* showed a weak phenotype with slightly increased nisin resistance but no synergistic effect was observed in the double deletion mutant CR099 *ΔΔporAH*. Interestingly, the triple deletion mutant *C. glutamicum* CR099 *ΔΔΔporABH* showed an more pronounced phenotype as the two single deletion mutants CR099 *ΔporA* and *CR099 ΔporABH* with approx. 2-fold higher resistance against nisin compared to the wildtype (**Figure 5A**). This indicates that nisin possibly translocate across mycolic acid porins and deletion of a single porin may be compensated by the presence of the remaining porins.

**Figure 5:**
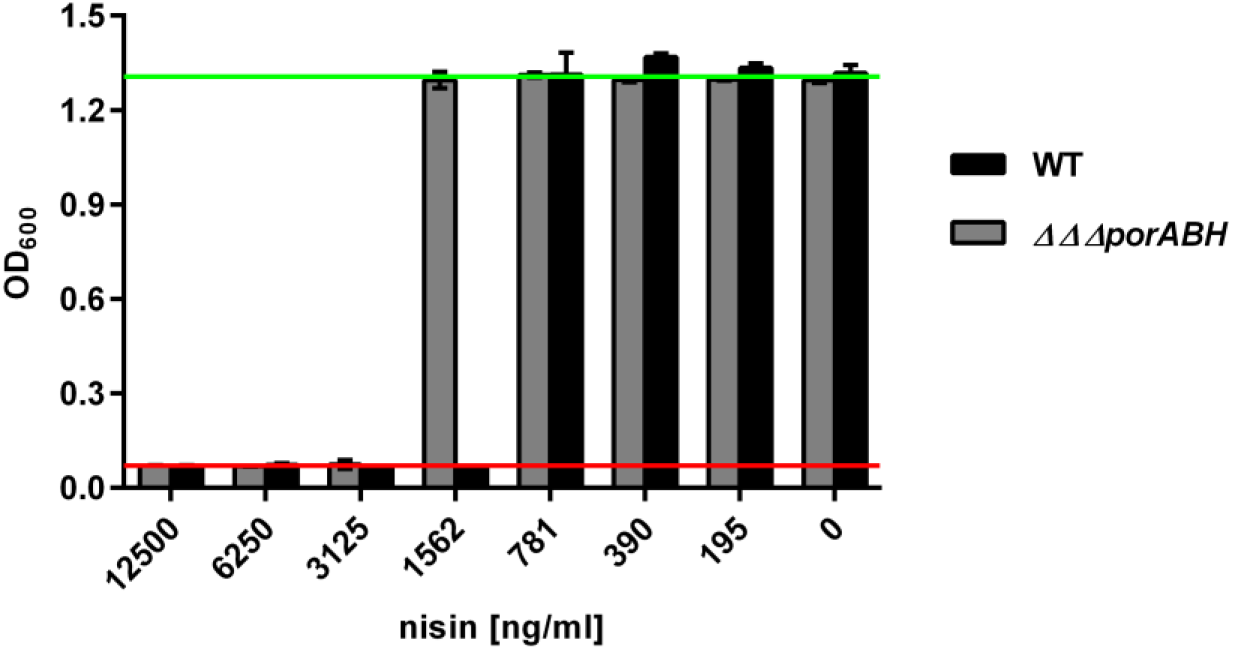
Resistance of *C. glutamicum* CR099 to *ΔΔΔporABH* nisin. Bacteria were grown in 96-well microtiter plates in 2xTY medium supplemented nisin at the indicated concentrations. OD_600_ was determined after 24 h of incubation. The red and green lines indicate OD_600_ of the positive (i.e. complete inhibition of growth) or negative (i.e. in the absence of nisin) control, respectively. Values are mean ± standard deviation of n = 6 cultivations of the test strain.

It remains to be elucidated if resistance to nisin may be further increased by additional mutations e.g. deletion of *porC*. Recently, a further potential porin encoded by *protX* (annotated as a protein with unknown function) was shown to share similarities to known porins and is also mycoloylated (Issa et al., 2017) and its deletion may increase nisin resistance. Moreover, the mycolic acid layer is not the only component which determines cell wall permeability as *C. glutamicum* cells are not completely covered with mycolic acids (Puech et al., 2001). To examine the impact of the mycolic acid membrane layer for nisin translocation, experiments with the *C. glutamicum* ATCC13032 *ΔotsAΔtreSΔtreY* that is unable to form mycolic acid layer when cultivated on fructose (Wolf et al., 2003) may be performed.

## 4 Conclusions

In the presented study, we aimed at improving resistance of *Corynebacterium glutamicum* to nisin by various approaches to facilitate heterologous production of this antimicrobial peptide. Some of these approaches resulted in slightly increased resistance (e.g. expression of the Bce-type ABC transporter VraDE from *S. aureus*, supplementation of growth media with CaCl_2_ or deletion of multiple porins). By contrast, other approaches known to increase nisin resistance of other organisms did not show any effect. These results suggest that transfer of nisin resistance mechanisms of other organisms to *C. glutamicum* is not trivial and does not result in dramatic improvements of resistance. Although there may be room for improvement, e.g. by combining approaches that have allowed an increase in resistance here, it is quite possible that levels of nisin resistance required to produce nisin at levels comparable to natural producers may not be achieved.

## 6 CRediT authorship contribution statement

D. Weixler: Conceptualization, Methodology, Investigation, Validation, Formal analysis, Writing - original draft, Writing - review & editing.

O. Goldbeck: Investigation, Writing - review & editing.

B.J. Eikmanns: Writing - review & editing.

G.M. Seibold: Conceptualization, Methodology, review & editing.

C.U. Riedel: Conceptualization, Methodology, Validation, Formal analysis, Writing - original draft, Writing - review & editing.

## 8 Acknowledgements

This project has received funding from the Bio Based Industries Joint Undertaking under the European Union’s Horizon 2020 research and innovation program (grant agreement No 790507). The authors gratefully acknowledge support by Karina Haas in cloning deletion constructs.

