## Supplementary data for "Towards improved resistance of *Corynebacterium glutamicum* against nisin"

ORCID: 0000-0001-7134-7085

16 **Table S1:** Bacterial strains and plasmids used in this study.

| Strain | Relevant characteristics | Source |
|---|---|---|
| ***Escherichia coli*** | | |
| DH5α | cloning host | (Hanahan, 1983) |
| ***Lactococcus lactis subsp. lactis*** | | |
| B1629 | nisin Z producer | Collection of Dzung Diep (unpublished) |
| ***Staphylococcus aureus*** | | |
| ATCC 43300 | MRSA type strain, methicillin-resistant | ATCC |
| ***Corynebacterium lactis*** | | |
| RW3-42 | type strain | (Wiertz et al., 2013) |
| ***Corynebacterium glutamicum*** | | |
| CR099 | *C. glutamicum* ATCC 13032 ΔCGP1 ΔCGP2 ΔCGP3 ΔISCg1 ΔISCg2; cured of prophages CGP1, CGP2 and CGP3 and insertion elements ISCg1 and ISCg2 | (Baumgart et al., 2013) |
| ΔporA | Deletion of porin coding gene *porA* (cg3008) | this study |
| ΔporH | Deletion of porin coding gene *porH* (cg3009) | this study |
| ΔporB | Deletion of porin coding gene *porB* (cg1109) | this study |
| ΔΔporAH | Deletion of porin coding genes *porA* and *porH* | this study |
| ΔΔΔporAHB | Deletion of porin coding genes *porA, porH* and *porB* | this study |
| **Plasmid** | **Relevant characteristics** | **Source** |
| pEKEx2 | *E. coli*/*C. glutamicum* shuttle vector; P*tacI*; *lacI*$^{q}$; *oriC.g* from pBL1.; *oriE.c.* ColE1 from pUC18; Kan$^{r}$. | (Eikmanns et al., 1994) |
| pEKEx2-*nisI* | pEKEx2 derivative for expression of *nisI* from *L. lactis* B1629 | this study |
| pEKEx2-*nisFEG* | pEKEx2 derivative for expression of *nis F, nisE, nisG* from *L.lactis* B1629 | this study |
| pEKEx2-*nisI*$^{CO}$ | pEKEx2 derivative for expression of *nisI*$^{CO}$ synthesized for expression in *C. glutamicum* | this study |
| pEKEx2-*vraDE* | pEKEx2 derivative for expression of *vraD, vraE* from *S. aureus* ATCC 43300 | this study |
| pEKEx2-*cg2812-11* | pEKEx2 derivative for over-expression of *cg2812-11* from *C. glutamicum* | this study |
| pEKEx2-*cg3322-20* | pEKEx2 derivative for over-expression of *cg3322-20* from *C. glutamicum* | this study |
| pEKEx2-*cg1103* | pEKEx2 derivative for over-expression of *cg1103* from *C. glutamicum* | this study |
| pOGOduet | dual expression vector; ptac ptet; lacI, tetR; KanR | (Goldbeck and Seibold, 2018) |
| pOGOduet-*nisI-nisFEG* | dual expression of *nisI* (P*tet* controlled) and *nisFEG* ($P_{tac}$ controlled) from *L. lactis* B1629 | this study |

**Table S1:** continued.

| **Plasmid** | **Relevant characteristics** | **Source** |
|---|---|---|
| pJYS3Δ*crtYf* | expression vector for deletion of *crtYf* in *C. glutamicum* | (Jiang et al., 2017) |
| pJYS3-KH | deletion vector; constitutive expression of *cpF1*; KanR | this study |
| pJYS_sgdporA_up_do | pJYS derivative for expression of protospacer region and up/downstream flanking regions for porA deletion | this study |
| pJYS_sgdporH_up_do (WT) | pJYS derivative for expression of protospacer region and up/downstream flanking regions for porH deletion | this study |
| pJYS_sgdporH_up_do (ΔporA) | pJYS derivative for expression of protospacer region and flanking regions for porH deletion in ΔporA mutant | this study |
| pJYS_sgdporB_up_do | pJYS derivative for expression of protospacer region and up/downstream flanking regions for porB deletion | this study |

**Table S2:** Oligonucleotide primers and synthetic gene sequences used in this study.

| Primer | Sequence (5' → 3') | Purpose |
|---|---|---|
| px2_fw<br>px2_rv | CACTCCCGTTCTGGATAATG<br>GCTACGGCGTTTCACTTCTG | Control primer flanking MCS pEKEx2 and pXMJ19 |
| cg2812 fw<br>cg2812 rv | CCTGCAGGTCGACTCTAGAGGCTAGCAAGGAGTTTTCATGAGTAACCCTGCCGCG<br>GAATTCATGAAAACTCCTTTTAGGCAATATCCTCAATTCCGTTC | Amplification and assembly of *cg2812* in px2 |
| cg2811 fw<br>cg2811 rv | TATTGCCTAAAAGGAGTTTTCATGAATTCCGGTTCCACAATG<br>ATTCGAGCTCGGTACCCGGGCCATGGCTAGTCGGTAATCGCATC | Amplification and assembly of *cg2811* in px2 |
| cg3322 fw<br>cg3322 rv | CCTGCAGGTCGACTCTAGAGGGTACCAAGGAGTTTTCTTGGCCCCGAAGAAATTAATC<br>GGCTCATGAAAACTCCTTCTAAATCACCTGGCCCAC | Amplification and assembly of *cg3322* in px2 |
| cg3321 fw<br>cg3321 rv | GGTGATTTAGAAGGAGTTTTCATGAGCCTCATCGAAATG<br>GGCTCATGAAAACTCCTTTCATGAGTGTTTCACCTC | Amplification and assembly of *cg3321* in px2 |
| cg3320 fw<br>cg3320 rv | ACACTCATGAAAGGAGTTTTCATGAGCCTTGCAGAATCAATTC<br>ATTCGAGCTCGGTACCCGGGGAGCTCTTACTCATAACGCAAGGC | Amplification and assembly of *cg3320* in px2 |
| cg2812-11 intseq1<br>cg2812-11 intseq2 | ATCCCACCATCGAGGAAATC<br>CTCTTCGGCTCTGCTCTTGG | Sequencing primer binding *cg2812-11* internally |
| cg3322-20 intseq1<br>cg3322-20 intseq2<br>cg3322-20 intseq3 | GGCCTGGAACAATCAATTGC<br>CAGCGACGACAACAAAGTAG<br>AACTGGTGCGTTGGATTCTG | Sequencing primer binding *cg3322-20* internally |
| vraD fw<br>vraD rv | CCTGCAGGTCGACTCTAGAGGTCGACAAGGAGTTTTCATGACAATATTATCAGTGCAAC<br>AATGTCATGAAAACTCCTTTTAAATGTCATTTGAGACACC | Amplification and assembly of *vraD* in px2 |
| vraE fw<br>vraE rv | TGACATTTAAAAGGAGTTTTCATGACATTTAACCATATCGTTTTC<br>ATTCGAGCTCGGTACCCGGGGAGCTCTTAAATGGTTTTCTTAATCAATTTG | Amplification and assembly of *vraE* in px2 |
| cg1103 fw<br>cg1103 rv | CCTGCAGGTCGACTCTAGAGAAGGAGTTTTCATGAAAGACGCTTCACAGTCC<br>ATTCGAGCTCGGTACCCGGGCTAGCTGTGGCTTGGGGC | Amplification and assembly of *cg1103* in px2 |
| cg1103seq1<br>cg1103seq2<br>cg1103seq3 | AGTGCTTGCTGCGCTTAATC<br>GTTGCGTGGTATTCCGTTTG<br>AGTTTCCGCTGCCGAATTGG | Sequencing primer binding *cg1103* internally |
| iPCR fw<br>iPCR rv | ℗-ACCCACGGGCCCGGTGAACAGTTG<br>℗-GATTGACAGCTAGCTCAGTCCTAGG | Amplification of pJYSΔ*crtYf* backbone and removal of flanking regions |
| Spacer fw<br>Sapcer rv | GATAATTTAAATCCTCGTCGTTGCTCCTCAG<br>GATAGGATCCGCAAAGCAAGACGCCCGGTTTG | Amplification of 500 bp non-codung fragment |
| upsite_porA fwd<br>upsite_porA rv | GCTAGCTGTCAATCTAGC CCATCTAACATTTCTGCAGG<br>CAAGCAGACCGTAAACGTTTTCCATTTTAAATTC | Amplification and assembly of *porA* upstream flanking region in pJYS_sgd*porA* |

**Table S2:** continued.

| Primer | Sequence (5’ → 3’) | Purpose |
|---|---|---|
| dosite_porA fwd<br>dosite_porA rv | AAACGTTTACGGTCTGCTTGGCTAATTAACTTC<br>TCACCGGGCCCTCTAGACCCAACAGTACGGGCACCACG | Amplification and assembly of *porA* downstream flanking region in pJYS_sgd*porA* |
| gnΔporA_fwd<br>gnΔporA_rv | GTAGTGTTACACCTTCATGGGG<br>GGTGGATTCGCGTGGTGATGG | Sequencing primer binding on *C. glutamicum* genome flanking *porA* deletion site |
| up_porH fwd<br>up_porH rv | GCTAGCTGTCAATCTAGCCCAAGATATTGCTTTTCGACG<br>TCTCTTAGGAATCCATGAGAAATCTCCTTG | Amplification and assembly of *porH* upstream flanking region |
| doWT_porH fwd<br>doWT_porH rv | TCTCATGGATTCCTAAGAGAAATCCGATTTGGC<br>TCACCGGGCCCTCTAGACCCAAGGGAAGTCTTCGCGCC | Amplification and assembly of porH downstream region in pJYS_sgd*porH* for deletion in WT |
| doΔ_dporH fwd<br>doΔ_dporH rv | TCTCATGGATTCCTAAGAGAAATCCGATTTGGCTGATTG<br>TCACCGGGCCCTCTAGACCCTGATGTGGCGCTGGCCAG | Amplification and assembly of porH downstream region in pJYS_sgd*porH* for deletion in *ΔporA* |
| gnΔporH fwd<br>gnΔporH rv | ATTCGCGCACCTCAATTGCC<br>AACAGTACGGGCACCACGAG | Sequencing primer binding on *C. glutamicum* genome flanking *porH* deletion site |
| up_porB fwd<br>up_porB rv | GCTAGCTGTCAATCTAGCCCTCTGTATCAATTTGCGGAAC<br>CCTTTTAGGACTTCATGATTTTTAGGGCTC | Amplification and assembly of *porB* upstream flanking region |
| do_porB fwd<br>do_porB rv | AATCATGAAGTCCTAAAAGGTTCGGGGG<br>TCACCGGGCCCTCTAGACCCGGAAGAAGATAGGTTAGAGGAC | Amplification and assembly of *porB* downstream region in pJYS_sgd*porB* |
| gnΔporB fwd<br>gnΔporB rv | GCAATCGCTTGAGCGGACAC<br>GGCTCAAGGAAAAGCCCAAG | Sequencing primer binding on *C. glutamicum* genome flanking *porB* deletion site |
| **Gene** | **Sequence (5’→ 3’)** | **Size [bp]** |
| $nisI^{CO}$ | ATGCGCAAGTATCTGATCTTGATCGTAGCGCTGATTGGAATCACCGGACTTTCAGGGT<br>GCTATCAGACTAGCCAGAAGAAAGTGCGCTTTGACGAAGGCTCCTATACCAACTTCA<br>TCTTCGACAACAAGTCCTACTTTGTCACCGACAAGGAGATTCCGCAAGAGAATGTCA<br>ACAACTCGAAAGTGAAGTTCTACAACCTCCTGATTGTGGACATGAAGTCCGAAAAGC<br>TGCTCTCCTCCTCCAATAAGAACTCCGTAACGTTGGTCCTGAACAACATCTACGAAGC<br>CTCAGACAAATCGCTCTGTATGGGCATCAATGATCGGTACTACAAGATTCTGCCTGA<br>GTCGGACAAAGGTGCAGTCAAGGCTTTGCGTCTGCAGAACTTCGATGTGACCTCTGA<br>CATTTCGGATGACAACTTCGTGATTGGCAAGAACGATAGCCGCAAAATCGACTACAT<br>GGGTAACATCTACTCTATCTCCGATACCACGGTTTCAGATGAGGAACTTGGCGAATAT<br>CAGGATTTCCTTTCCGAAGTTCGCGTTTTCGATAGCGTTAGCGGTAAGTCCATTCCAC<br>GCTCAGAATGGGGTCGAATCGATAAAGACGGCTCCAATTCCAAGCAATCTCGTACAG<br>AGTGGGATTACGGTGAGATCCACTCTATCCGTGGAAAGTCTCTGACTGAAGCCTTTGC<br>AGTTGAGATCAATGACGATTTCAAACTCGCTACCAAAGTCGGCAACTAA | 738 |

$^{Cg}$: codon-optimized for *C. glutamicum*

**Table S3:** Oligonucleotides used for gene deletions. Protospacer sequences (5´->3´) are highlighted in bold letters

| Oligonucleotide | Sequence 5´ → 3´ | Purpose |
|---|---|---|
| OE_sg_univ. fwd | GGGCTAGATTGACAGCTAGCTCAGTCCTAGGTATAATGGATCCGAATTTCTACTGTTGTAGAT | Universal fwd OE-PCR primer for protospacer region product |
| OE_sg_Δ*porA* rv | CTGAGCCTTTCGTTTTATTTAAAT**TAGCCGATGAGGCCGGAGCCG**ATCTACAACAGTAGAAATTC | Specific rv OE-PCR primer for *porA* targeting protospacer |
| OE_sg_Δ*porH* rv | CTGAGCCTTTCGTTTTATTTAAAT**CCGAGGGTTTCCTTGAGAAGG**ATCTACAACAGTAGAAATTC | Specific rv OE-PCR primer for *porH* targeting protospacer |
| OE_sg_Δ*porB* rv | CTGAGCCTTTCGTTTTATTTAAAT**GCCATTGCTGCGATGCGGTGT**ATCTACAACAGTAGAAATTC | Specific rv OE-PCR primer for *porB* targeting protospacer |

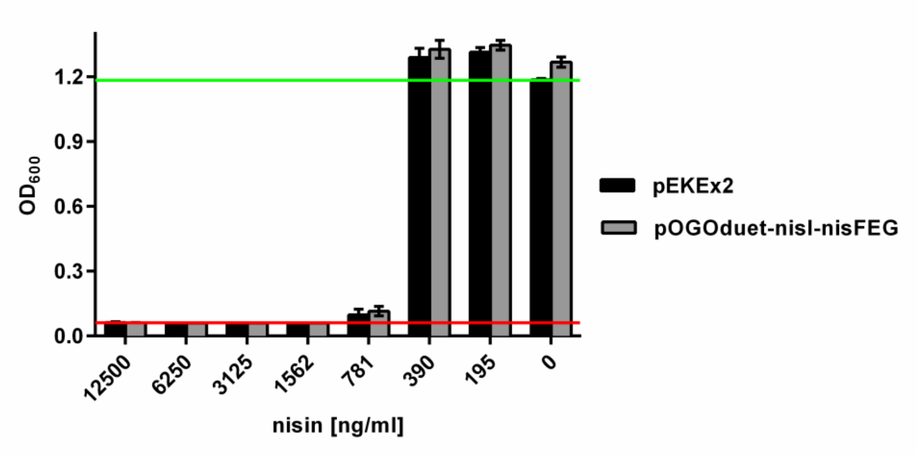


**Supplementary Figure S1: Resistance of *C. glutamicum* CR099/pOGOduet-*nisI-nisFEG* (grey bars) or the empty vector control strain CR099/pEKEx2 (black bars) to nisin.** Bacteria were cultivated in 2xTY medium in 96-well microtiter plates in the presence of nisin at the indicated concentrations. For induction of gene expression 0.1 mM IPTG was added. Optical density at 600 nm ($OD_{600}$) was determined after 24 h of incubation. The red and green lines indicate $OD_{600}$ of the positive (i.e. complete inhibition of growth) or negative (i.e. in the absence of nisin) control, respectively. All values are mean ± standard deviation of n = 3 cultivations of the test strain.
